# Interferon-induced transmembrane protein 3 (IFITM3) limits lethality of SARS-CoV-2 in mice

**DOI:** 10.1101/2021.12.22.473914

**Authors:** Ashley Zani, Adam D. Kenney, Jeffrey Kawahara, Adrian C. Eddy, Xiao-Liang Wang, Mahesh KC, Mijia Lu, Emily A. Hemann, Jianrong Li, Mark E. Peeples, Luanne Hall-Stoodley, Adriana Forero, Chuanxi Cai, Jianjie Ma, Jacob S. Yount

## Abstract

Interferon-induced transmembrane protein 3 (IFITM3) is a host antiviral protein that alters cell membranes to block fusion of viruses. Published reports have identified conflicting pro- and antiviral effects of IFITM3 on SARS-CoV-2 in cultured cells, and its impact on viral pathogenesis *in vivo* remains unclear. Here, we show that IFITM3 knockout (KO) mice infected with mouse-adapted SARS-CoV-2 experienced extreme weight loss and lethality, while wild type (WT) mice lost minimal weight and recovered. KO mice had higher lung viral titers and increases in lung inflammatory cytokine levels, CD45-positive immune cell infiltration, and histopathology, compared to WT mice. Mechanistically, we observed disseminated viral antigen staining throughout the lung tissue and pulmonary vasculature in KO mice, while staining was observed in confined regions in WT lungs. Global transcriptomic analysis of infected lungs identified upregulation of gene signatures associated with interferons, inflammation, and angiogenesis in KO versus WT animals, highlighting changes in lung gene expression programs that precede severe lung pathology and fatality. Corroborating the protective effect of IFITM3 *in vivo*, K18-hACE2/IFITM3 KO mice infected with non-adapted SARS-CoV-2 showed enhanced, rapid weight loss and early death compared to control mice. Increased heart infection was observed in both mouse models in the absence of IFITM3, indicating that IFITM3 constrains extrapulmonary dissemination of SARS-CoV-2. Our results establish IFITM3 KO mice as a new animal model for studying severe SARS-CoV-2 infection of the lung and cardiovascular system, and overall demonstrate that IFITM3 is protective in SARS-CoV-2 infections of mice.

## INTRODUCTION

(IFITM3) is an antiviral restriction factor that inhibits virus fusion with host cell membranes^1–3^, and potently blocks infection by numerous enveloped viruses of human health concern, including influenza, dengue, and Zika viruses^4^. This set of viruses primarily utilizes endocytic pathways for entry into cells, and fuses with endosome membranes where IFITM3 is abundantly localized and poised to restrict infection^5,6^. In contrast, viruses that fuse primarily at the plasma membrane, such as Sendai virus, are minimally affected by IFITM3^7,8^. Viruses that are able to fuse at both the plasma membrane or within endosomes, such as human metapneumovirus, can be partially restricted by IFITM3, with the portion of virus that fuses at the plasma membrane largely evading restriction^9^. These patterns of restriction hold true for the vast majority of the dozens of viruses that have been tested for IFITM3 inhibition, with the notable exception of the common cold coronavirus OC43, which utilizes dual cell entry pathways similarly to metapneumovirus, but showed enhanced, rather than reduced, infection when IFITM3 was expressed in target cells^10^. This unusual finding suggests that IFITM3 may have unanticipated effects on some coronaviruses, leading several groups to examine the impact of IFITM3 and related proteins on SARS-CoV-2, which remains a critical threat to worldwide human health.

SARS-CoV-2 uses cell surface and endosomal fusion strategies for infection^11,12^. Experiments studying effects of IFITM3 on SARS-CoV-2 infections identified opposing activities in which IFITM3 inhibits virus entry at endosomes as expected, but enhances virus entry at the plasma membrane^13^. The overall effects of IFITM3 remain controversial, with some reports indicating that IFITM3 primarily restricts cellular SARS-CoV-2 infection^13,14^, while others conclude that IFITM3 is a net enhancer of infection, particularly in lung cells^15–17^. Further, IFITM3 has been shown inhibit fusion of SARS-CoV-2 infected cells with neighboring cells (syncytia formation), which may be an important mechanism of viral spread within tissues^18,19^. The overall balance and relevance of the effects of IFITM3 in SARS-CoV-2 infection of cells and pathogenesis *in vivo* remain unclear.

We previously generated IFITM3 knockout (KO) mice to investigate *in vivo* effects of this antiviral restriction factor and confirmed a profound susceptibility of these mice to severe influenza virus infections^20^. These results were in agreement with studies linking deleterious single nucleotide polymorphisms in the human *IFITM3* gene to severe influenza^21–23^. Although some studies have suggested that *IFITM3* gene polymorphisms are risk factors for severe SARS-CoV-2 infection^24–28^, *IFITM3* variants did not emerge as top candidates in other large genome-wide SARS-CoV-2 susceptibility studies^29–31^. In addition to possible direct effects of *IFITM3* gene polymorphisms, defects in interferon responses that induce IFITM3 during infection have been linked to roughly 20% of severe COVID-19 cases, providing additional circumstances where IFITM3 insufficiency may be impacting disease^32,33^. Given the conflicting reports of IFITM3 activity on SARS-CoV-2 *in vitro*, coupled with the potential, but uncertain, role of IFITM3 in modulating SARS-CoV-2 disease severity, we investigated roles of IFITM3 in SARS-CoV-2 infection in the mouse model.

## RESULTS AND DISCUSSION

To determine the impact of IFITM3 on SARS-CoV-2 disease severity, we infected wild type (WT) C57BL/6 mice and corresponding IFITM3 KO animals with the mouse-adapted SARS-CoV-2 strain MA10^34^. Importantly, our virus stock was generated with a stringent plaque purification and sequencing protocol, allowing us to utilize virus lacking commonly observed tissue culture adaptations that attenuate pathogenicity^35^. After infection, we observed a greater than 10% weight loss in WT mice with recovery to full weight by day 8 post infection (**Fig 1A**). In contrast, IFITM3 KO mice lost significantly more weight starting at day 1 post infection. This accelerated weight loss continued through day 5 post infection, at which point all IFITM3 KOs had either died or met humane endpoint criteria due to severe illness and lack of movement (**Fig 1A**). We conclude that IFITM3 provides protection against severe, lethal SARS-CoV-2-induced disease in mice.

**Figure 1:**
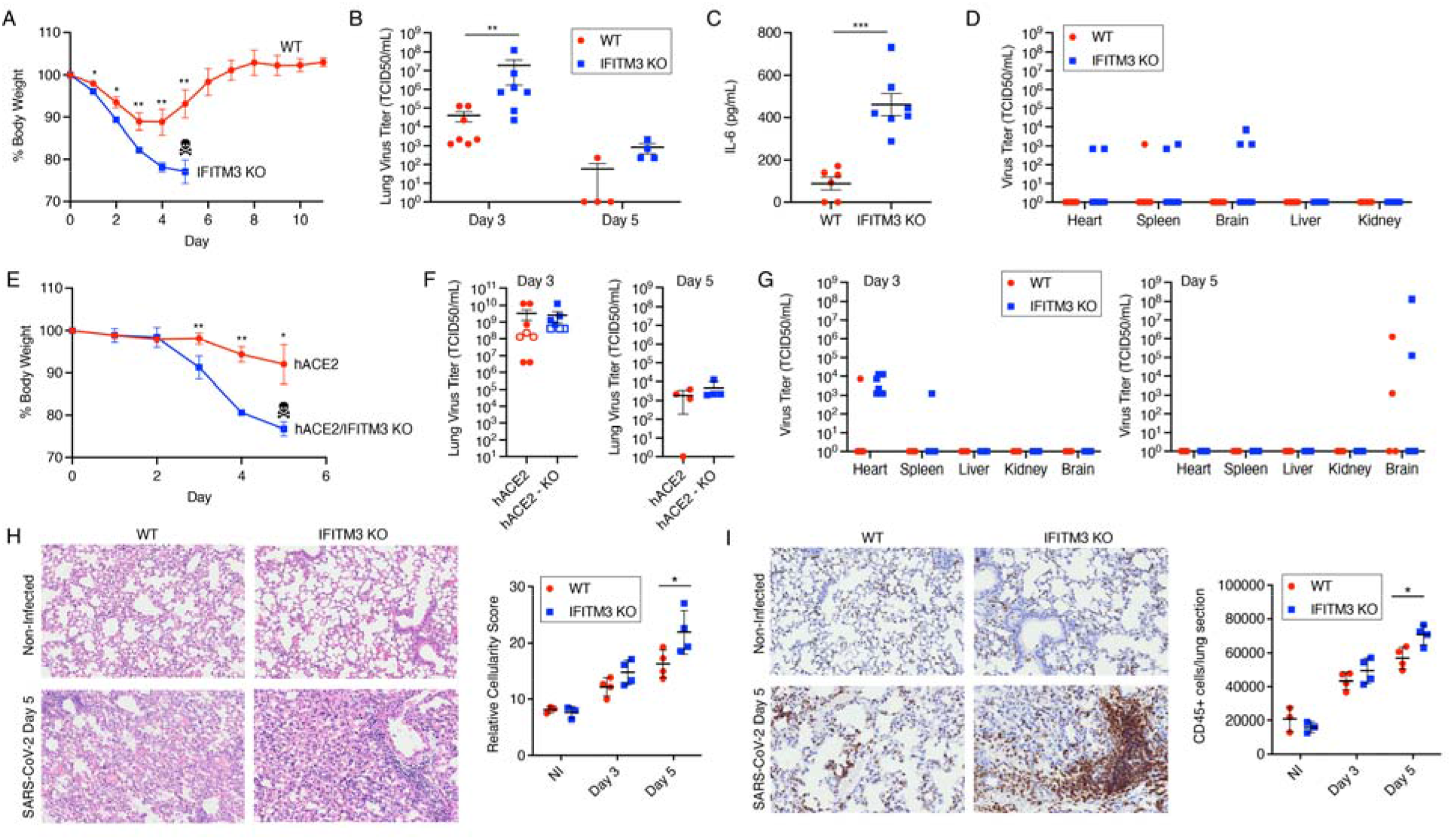
IFITM3 limits the severity of SARS-CoV-2 infections. **A-D)** WT and IFITM3 KO mice were intranasally infected with 10^5^ TCID50 SARS-CoV-2 MA10. **A)** Weight loss (*p < 0.05, ** p < 0.01, ANOVA with Bonferoni’s multiple comparison’s test, data from three independent experiments, scull and crossbones indicates that KO mice did not survive beyond this timepoint), **B)** viral titers in lung homogenates at the indicated days post infection (**p < 0.01, Mann-Whitney test, data from two independent experiments), **C)** IL-6 levels in lung homogenates on day 3 post infection (***p < 0.001, t-test, data from two independent experiments), and **D)** viral titers in extrapulmonary organs on day 3 post infection were measured. **E-G)** K18-hACE2 (hACE2) and K18-hACE2/IFITM3 KO mice were infected with 10^4^ TCID SARS-CoV-2 WA1. **E)** Weight loss (*p < 0.05, ** p < 0.01, ANOVA with Bonferoni’s multiple comparison’s test, scull and crossbones indicates that KO mice did not survive beyond this timepoint), **F**) viral titers in lung homogenates at the indicated days post infection (data for day 3 are from two independent experiments, data points represented by open circles/squares are from a single experiment in which **p < 0.01, Mann-Whitney test), and **G)** viral titers in extrapulmonary organs on the indicated days post infection were measured (extrapulmonary organ data are from one experiment, except for the heart, which are from two independent experiments). **H,I)** WT and IFITM3 KO mice were intranasally infected with 10^5^ TCID50 SARS-CoV-2 MA10 or mock infected with PBS (Non-Infected, NI). Representative lung images from mice at the indicated times post infection were stained with hematoxylin and eosin **(H)** or anti-CD45 **(I)**. Example images and staining quantifications from multiple mice are shown (*p < 0.05, ANOVA with Bonferoni’s multiple comparison’s test). For all graphs of virus titers, data points at the x axis represent samples below the limit of detection.

We next measured viral titers at days 3 and 5 post infection, and found that IFITM3 KO lung titers were significantly higher than WT titers on day 3 (**Fig 1B**). Consistent with higher virus measurements, elevated levels of the inflammatory cytokine IL-6 were also detected in KO versus WT lungs at day 3 post infection (**Fig 1C**). Viral titers were decreased in both groups at day 5 compared to day 3 (**Fig 1B**). Nonetheless, complete virus clearance was delayed in IFITM3 KOs, with all KO mouse lungs remaining positive for virus at day 5, while only one of four WT lungs were positive for live virus at this timepoint (**Fig 1B**). At day 3 post infection, we also detected a low level of live virus in the heart, spleen, and brain in a subset of the KO animals, while no virus was detected in liver or kidney tissue (**Fig 1D**). Live virus was not detected in any of these extrapulmonary organs in WT or KO animals at day 5 post infection (not shown). In sum, we observed that the enhanced illness severity in IFITM3 KO mice was accompanied by increased viral titers and delayed clearance.

Mouse-adapted SARS-CoV-2 differs from the parental human isolate by only seven amino acids^34^ and utilizes endogenously expressed murine ACE2 as the virus receptor for infection, making it a highly relevant model for studying viral tropism and pathogenesis *in vivo*. Nonetheless, we also sought to examine the *in vivo* role of IFITM3 in infection with WA1, a nonadapted SARS-CoV-2 strain isolated from humans, which we also stringently propagated and sequenced. For this, we utilized mice expressing human (h)ACE2 under control of the K18 keratin promoter (K18-hACE2 mice) crossed with IFITM3 KOs. We observed that K18-hACE2/IFITM3 KO mice lost significantly more weight than K18-hACE2 control mice, and that all of the K18-hACE2/IFITM3 KO mice succumbed to infection by day 5 (**Fig 1E**), at which point we ended our experiments. High virus titers at day 3 post infection were detected in both control and KO mice, with one of two experiments showing significantly higher titers in IFITM3 KOs and a second experiment showing a trend toward higher titers in the KOs (**Fig 1F**). Further, cardiac dissemination of virus was detected in 8 out of 8 IFITM3 KOs compared to 1 out of 8 control mice (**Fig 1G**), confirming that IFITM3 restricts cardiac dissemination of SARS-CoV-2. Overall, a protective effect of IFITM3 in SARS-CoV-2 infections was identified using both mouse-adapted and human isolates.

To mechanistically characterize roles of IFITM3 in limiting SARS-CoV-2 lung pathogenesis, we performed hematoxylin and eosin staining on lung sections from WT and IFITM3 KO animals after infection with SARS-CoV-2 MA10. All lungs from infected animals showed areas of consolidated tissue, cellular infiltration, and inflammation (**Fig 1H**). We observed that larger portions of the lungs were afflicted in IFITM3 KO samples, and thus quantified cellularity scores to measure loss of open airspace for individual sections from infected WT and IFITM3 KO mouse lung sections. This unbiased method confirmed increased pathology in infected IFITM3 KO versus WT lungs at day 5 post infection (**Fig 1H**), correlating with the increased illness and viral titers that we observed in these animals (**Fig 1A, B**). This conclusion was supported further by immunohistochemical staining and quantification of CD45-positive immune cell infiltration into the lung, which was elevated in IFITM3 KO versus WT mice at day 5 (**Fig 1I**). Thus, lung pathology resulting from SARS-CoV-2 infection was exacerbated in IFITM3 KO mice.

Given that IFITM3 generally blocks infection and spread of certain viruses, we investigated the effects of IFITM3 on viral infection patterns in the lungs by staining for viral antigen (N protein) in WT and IFITM3 KO lung sections at day 2 post infection with SARS-CoV-2 MA10, representing an early timepoint at which viral titers are high^34^. We detected robust staining of the airways in both groups, but observed more disseminated staining throughout IFITM3 KO lungs (**Fig 2A, B**). Indeed, large portions of WT lungs showed no staining for viral antigen, while virus was detected throughout KO lungs (**Fig 2A,B**). Further, accumulation of viral antigen that likely represents shedding of necrotic, highly infected cells into the bronchioles was observed in KO lungs, even at this early timepoint (**Fig 2B**). Infected cells associated with blood vessels could be seen in WT lungs primarily when adjacent to highly infected airways (**Fig 2A**), while blood vessels associated with infected cells were readily apparent throughout IFITM3 KO lungs (**Fig 2B**). Quantification of viral staining in lung sections from multiple mice confirmed the significant increase in viral antigen staining in IFITM3 KO versus WT mice (**Fig 2C**). Overall, viral antigen imaging confirmed increased viral titers and pathology (**Fig 1B, H**), but also revealed diffuse infection of cells, including the vasculature, in IFITM3 KO lungs.

**Figure 2:**
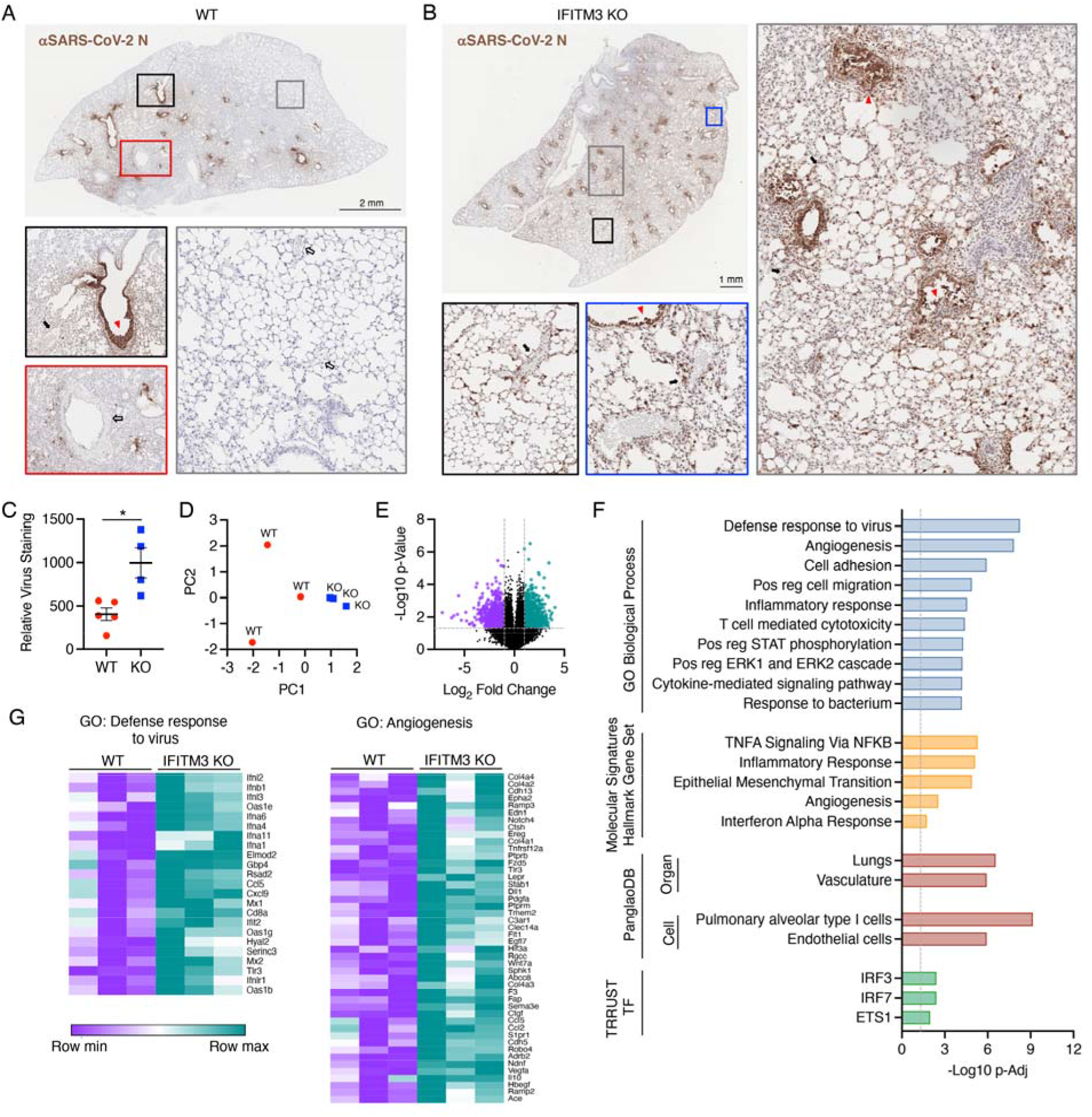
Diffuse SARS-CoV-2 infection of IFITM3 KO versus WT lungs induces increased inflammatory and angiogenic gene expression programs. WT and IFITM3 KO mice were intranasally infected with 10^5^ TCID50 SARS-CoV-2 MA10. **A,B)** Representative lung sections from **A)** WT or **B)** IFITM3 KO animals stained for SARS-CoV-2 N protein. Colored boxes overlaying the full lung images correspond to the displayed magnified regions. Red triangles, infected airways; solid black arrow, non-infected blood vessels; open black arrow, infected blood vessels. **C)** Quantification of lung section staining for multiple mice as depicted in **A** and **B** (*p < 0.05, t-test). **D-G)** RNA sequencing analysis of RNA extracted from infected WT and IFITM3 KO lungs on day 2 post infection. **D)** Principal components analysis comparing WT and KO RNA samples. **E)** Volcano plot of differential gene expression comparing IFITM3 KO versus WT. Purple, 958 genes downregulated in KO versus WT; Green, 907 genes upregulated in KO versus WT. **F)** Upregulated genes in KO versus WT were subjected to pathway and ontology analysis. The top ten most significant GO Biological Processes are shown (-Log10 p-Elim pruning value is graphed), the top five most significant Hallmark gene sets, the significant associations from PanglaoDB analysis, and the significant associations from TRRUST transcription factor analysis are shown. **G)** Heat maps for genes from the top two most significant GO Biological Processes as shown in **F**.

We next examined the global impact of loss of IFITM3 on transcriptional programs at day 2 post infection with SARS-CoV-2 MA10 by performing lung RNA sequencing. Dimensionality reduction approaches showed divergence of gene signatures in infected WT versus KO lungs (**Fig 2D**), corresponding to 1,865 differentially expressed genes between the groups (≥ |2| fold change, p-value < 0.05; a list of differentially expressed genes, expression values, fold changes, and p-values are shown in **Supplementary Table 1**). GO term enrichment analysis of genes upregulated in infected IFITM3 KO versus WT lungs revealed significant increases in antiviral defense responses, angiogenesis, and inflammation (**Fig 2F, Supplementary Table 2**). Among the antiviral and inflammatory genes increased in KO versus WT animals were multiple type I interferon genes, including *Ifnb1* and *Ifna4*, and canonical interferon-stimulated genes, such as *Oas1b*, *Mx1*, and *Ifit2* (**Fig 2G**). Expression of chemokine genes *Ccl5* and *Cxcl9* was also potentiated in KO lungs relative to WT (**Fig 2G**), consistent with the increased recruitment of CD45-positive immune cells that we observed (**Fig 1I**). Classic angiogenesis-driving genes with significantly elevated expression in KO versus WT animals included *Vegfa*, *Hif3a*, *Wnt7a*, and *Pdgfa* (**Fig 2G**). These statistically significant increases in activation of inflammatory antiviral and angiogenesis pathways in KO lungs were further confirmed by Molecular Signature Database Hallmark gene set enrichment analysis (**Fig 2F**). Likewise, analysis with the PanglaoDB database of single cell RNA sequencing experiments identified significant associations with gene signatures specific to pneumocytes and endothelial cells within the KO-specific upregulated genes (**Fig 2F**). Furthermore, TRRUST Transcriptional Regulatory Network database analysis identified interferon regulatory factors, IRF3 and IRF7, as well as the pro-angiogenic protein ETS1, as transcriptional factors likely involved in upregulation of genes in infected IFITM3 KO versus WT lungs (**Fig 2F**). Thus, multiple independent analyses of our data converged on inflammatory interferon and angiogenic/endothelial nodes as key pathways that precede the uniquely severe manifestations of SARS-CoV-2 infections occurring in the absence of IFITM3. Analysis of the downregulated genes in IFITM3 KO versus WT lungs indicated association with muscle gene signatures, possibly suggesting infection and elimination of lung smooth muscle cells by SARS-CoV-2 infection, or disruption of regenerative responses (**Supplementary Table 2**). Overall, our results demonstrate that IFITM3 restrains SARS-CoV-2 replication and spread through the lungs and cardiovascular system, thus limiting induction of pathological interferon and angiogenesis gene expression programs.

Increased induction of interferon and interferon-responsive genes may be the result of heightened levels of virus replication seen in IFITM3 KO animals (**Fig 1B** and **Fig 2A, B**). Alternatively, these results are consistent with feedback inhibition of type I interferon induction and inflammation by IFITM3, as has been previously suggested^36,37^. Type I interferons provide a double-edged sword in COVID-19, limiting virus replication early in infection while also contributing to pathological inflammation^38,39^. Furthermore, angiogenesis and thrombosis rapidly emerged as hallmarks of SARS-CoV-2-associated disease^40^. The observed upregulation of angiogenic gene programs may result from direct infection of blood vessel cells, or from infected cells mediating bystander signaling. We also noted that several coagulation-associated genes had enhanced upregulation in the KO animals, such as *Vwf*, *F2r*, *F2rl3*, *Plat*, *Fgg*, *F3*, and *Thbd* (**Supplementary Table 2**), consistent with exacerbated endothelial cell dysfunction occurring in the IFITM3 KO animals. Taken together, IFITM3 KO mice represent an important new tool for investigating pathological mechanisms active in severe SARS-CoV-2 infections, including angiogenic and pro-thrombotic signals. In addition to uncovering the utility of IFITM3 KO mice for future investigations of SARS-CoV-2 pathogenesis mechanisms, the results of our study demonstrate that IFITM3 restrains virus replication, limits activation of inflammatory and angiogenic hallmarks of severe disease, and overall limits pathology in SARS-CoV-2 infections *in vivo*.

## MATERIALS AND METHODS

### Biosafety

All experiments with SARS-CoV-2 were performed in the Ohio State University (OSU) Biosafety Level 3 (BSL3) facility. Experimental procedures were approved by the OSU Institutional Biosafety Committee and OSU BSL3 Advisory Group. Research samples analyzed outside of BSL3 containment were decontaminated according to experimentally validated and institutionally approved protocols.

### Virus stocks and titering

SARS-CoV-2 strain USA-WA1/2020 (WA1) and mouse-adapted SARS-CoV-2 strain MA10 (generated by the laboratory of Dr. Ralph Baric, University of North Carolina) were obtained from BEI Resources. Both viruses were plaque purified on Vero-TMPRSS2 cells (kindly provided by Dr. Shan-Lu Liu, The Ohio State University), and viral genomes from individual plaques were sequenced to identify virus lacking mutations in the furin cleavage site of the Spike protein. Virus plaques with intact furin cleavage sites were propagated on Vero-TRMPSS2 cells. Virus-containing supernatants were aliquoted and flash frozen in liquid nitrogen, and stored at −80C. Virus stocks were again sequenced to ensure the absence of furin cleavage site mutations. Viral stocks and research samples were titered via TCID50 measurements on Vero E6 cells using 10-fold serial dilutions in triplicate. Presence of virus replication in each well was assessed by visual examination of cytopathic effect along with antibody staining for N protein antigen (mouse monoclonal antibody 40143-MM08, Sino Biological) with fluorescent secondary antibody detection. TCID50 titers were calculated using the Reed-Muench method.

### Mice and Infections

IFITM3 KO mice with a 53 base pair deletion in exon 1 of the *Ifitm3* gene were described previously^20^. K18-hACE2 hemizygous mice were purchased from Jackson Laboratories. IFITM3-KO mice were crossed with K18-hACE2 mice to obtain F1 mice possessing a K18-hACE2 allele with heterozygous IFITM3 KO. F1 mice were then bred with IFITM3 KO mice to generate mice carrying hACE2 transgene and homozygous KO of IFITM3. Both male and female mice were used in our experiments. Mice were infected intranasally under anesthesia with isoflurane. Mouse organs were collected and homogenized in PBS for ELISAs or titering, homogenized in Trizol for extraction of RNA, or fixed in 10% formalin for histology.

### ELISAs and Histology

IL-6 ELISAs were performed on fluid from lung homogenates using an R&D Systems Duoset ELISA kit (catalogue # DY406) according to manufacturer’s instructions. For lung histology, lung tissue samples were fixed in formalin for 7 days at 4°C, followed by embedding in paraffin. For H&E and anti-CD45 staining, lungs were sectioned, stained, and imaged by the OSU Comparative Pathology and Mouse Phenotyping Shared Resource. Quantification of these images was performed using ImageJ software and the color deconvolution method. Staining for SARS-CoV-2 antigen was performed and imaged by Histowiz (Histowiz.com, Brooklyn, NY, USA).

### RNA sequencing and Informatics

RNA was extracted from lung tissue using TRIzol (Invitrogen). RNA library preparation and sequencing was performed by GENEWIZ, LLC./Azenta US, Inc (South Plainfield, NJ, USA) with the following methods as provided by GENEWIZ: “The RNA samples received were quantified using Qubit 2.0 Fluorometer (ThermoFisher Scientific, Waltham, MA, USA) and RNA integrity was checked using TapeStation (Agilent Technologies, Palo Alto, CA, USA). The RNA sequencing libraries were prepared using the NEBNext Ultra II RNA Library Prep Kit for Illumina using manufacturer’s instructions (New England Biolabs, Ipswich, MA, USA). Briefly, mRNAs were initially enriched with Oligod(T) beads. Enriched mRNAs were fragmented for 15 minutes at 94 °C. First strand and second strand cDNA were subsequently synthesized. cDNA fragments were end repaired and adenylated at 3’ends, and universal adapters were ligated to cDNA fragments, followed by index addition and library enrichment by PCR with limited cycles. The sequencing libraries were validated on the Agilent TapeStation (Agilent Technologies, Palo Alto, CA, USA), and quantified by using Qubit 2.0 Fluorometer (ThermoFisher Scientific, Waltham, MA, USA) as well as by quantitative PCR (KAPA Biosystems, Wilmington, MA, USA). The sequencing libraries were multiplexed and clustered onto a flowcell. After clustering, the flowcell was loaded onto the Illumina HiSeq instrument according to manufacturer’s instructions. The samples were sequenced using a 2×150bp Paired End (PE) configuration. Image analysis and base calling were conducted by the HiSeq Control Software (HCS). Raw sequence data (.bcl files) generated from Illumina HiSeq was converted into fastq files and de-multiplexed using Illumina bcl2fastq 2.17 software. One mis-match was allowed for index sequence identification.”

Resulting fastq files were analyzed by ROSALIND^®^ (https://rosalind.bio/), with a HyperScale architecture developed by ROSALIND, Inc. (San Diego, CA). Read Distribution percentages, violin plots, identity heatmaps, and sample MDS plots were generated as part of the QC step. Statistical analysis for differential gene expression was done using the “limma” R library^41^. The principal components analysis and volcano plots were re-formatted in PRISM graphing software using values downloaded from ROSALIND. Heatmaps were generated using gene expression values downloaded from ROSALIND and Morpheus software (https://software.broadinstitute.org/morpheus). Hypergeometric distribution was used to analyze the enrichment of pathways, gene ontology, and other ontologies. The topGO R library was used to determine local similarities and dependencies between GO terms in order to perform Elim pruning correction. Several database sources were referenced for enrichment analysis, including MSigDB^42,43^, TRRUST^44^, and PanglaoDB^45^. Enrichment was calculated relative to a set of background genes relevant for the experiment.

## ACKNOWLEDGMENTS

The authors thank Dr. Kara Corps (OSU Comparative Pathology and Mouse Phenotyping Core Facility) for overseeing lung sectioning and staining, and Dr. Eugene Oltz (OSU) for critical reading and editing of the manuscript. Research in the Yount laboratory is supported by NIH Grants AI130110, AI151230, HL154001, AI142256, and CA260582, and an American Lung Association COVID-19 and Emerging Respiratory Viruses Research Award. AZ is supported by an NSF-GRFP fellowship grant. MEP and MKC are supported by NIH grant U19AI42733.

